# Establishment and characterization of mammary organoids from non-traditional model organisms

**DOI:** 10.1101/2021.01.15.426833

**Authors:** Arianna P. Bartlett, Gerlinde R. Van de Walle

## Abstract

Mammary organoid (MaO) models are only available for a few traditional model organisms, limiting our ability to investigate mammary gland development and cancer across the diverse taxa of mammals. For example, horses are mammals with a similar mammary anatomy and function as humans, but they have a remarkably low incidence of mammary cancer, making the development of MaOs in non-traditional model organisms attractive, particularly in comparative cancer research. This study established equine mammary organoids (EqMaOs) from mammary gland tissue fragments and evaluated parameters including diameter, budding, and growth stage in non-budding EqMaOs, in cultures with increasing concentrations of epidermal growth factor (EGF), a key growth factor implicated in mammary gland development. Our findings showed that EqMaO diameter is not influenced by EGF concentration, whereas number of EqMaOs with budding and stage in non-budding EqMaOs are positively influenced by increasing EGF concentration. EqMaOs also formed protrusions with putative functions, including organoid fusion and sensory functions. We further characterized EqMaOs by the presence of myoepithelial and luminal cells using immunohistochemistry and used the hormone prolactin to stimulate milk secretion, as illustrated by β-lactoglobulin expression, in these EqMaOs. Additionally, we showed that our method to establish MaOs is widely applicable to additional non-traditional mammalian model organisms such as cat, pig, deer, rabbit, and prairie vole. Collectively, MaO models across species will be a useful tool for comparative developmental and cancer studies.

**Summary statement:** Mammary organoids can be established from various mammals by embedding mammary tissue fragments into a 3D matrix, providing a high-throughput, physiologically accurate model for comparative studies centered on mammary gland development and cancer.

## INTRODUCTION

The mammary gland arises from the ectoderm to form a rudimentary ductal tree which undergoes developmental expansion during puberty, where it forms a network of ducts and alveoli. Mammary development has been studied largely using rodent models, for which there are many tools available including transgenic mice and several well-established culture models (Darcy et al. 2000, Smits et al. 2007, Inman et al. 2015, Ngyuen-Ngoc et al. 2015). In contrast, our knowledge of mammary development and lactogenesis in other species, which we refer to as “non-traditional model organisms,” is somewhat limited, apart from studies in cows (Akers 2017, Geiger 2019) and pigs (Zhang et al. 2018, Hurley 2019), which are studied based on their importance in the dairy and meat industry. However, cell and tissue culture models from additional non-traditional species, both domesticated and wild, are lacking. Currently available models to study non-traditional model organisms are 3D suspension cultures called mammospheres (Rauner et al. 2018, Sumbal et al. 2020) and 2D adherent cultures established from these mammospheres, called mammosphere-derived epithelial cells (MDECs) (Spaas et al. 2012, Bussche et al. 2016, Ledet et al. 2018, Ledet et al. 2018, Ledet et al. 2020). Mammospheres are a culture of free-floating heterogenous mammary epithelial cell colonies enriched for stem/progenitor cells, which are generated using ultra-low attachment plates (Dontu et al. 2003). While these mammospheres have cellular heterogeneity and may be easily genetically manipulated, they lack *in vivo*-like mammary gland tissue structures and do not recapitulate cell-to-cell contact and signaling accurately (Srivastava et al. 2020). However, all these traits can be recapitulated in mammary organoids (MaOs), which are self-organizing tissues grown in a 3D matrix (Fatehullah et al. 2018, Sumbal et al. 2020, Srivastava et al. 2020). These MaOs can be established from several sources including primary explants, such as single cells or tissue fragments, reconstituted aggregates, suspension cultures, cell lines such as normal mammary epithelium or mammary cancer cell lines, mammary stem cells, and lastly, embryonic stem cells or induced-pluripotent stem cells (Sumbal et al. 2020, Fatehullah et al. 2018, Srivastava et al. 2020).

Currently, MaO models are well-established for mouse (Ngyuen-Ngoc et al. 2015), rat (Darcy et al. 2000), and human (Miller et al. 2017). MaO models for other species, however, are lacking with the exception of a few non-traditional model organisms including dog (Cocola et al. 2017), marmoset (Wu et al. 2016), and cow (Ellis 1998, Martignani et al. 2018). Establishing MaO models for additional non-traditional model organisms, present some challenges, for example, due to the lack of known markers to univocally identify progenitor/stem cells in the mammary gland of most non-traditional model organisms. Still, the availability of MaOs from non-traditional model organisms is important from the perspective of providing a physiologically relevant model for comparative mammary development. Although it is well-accepted that mammary development varies between species, most notably related to lactogenesis strategies, there are no current methods to perform multiple-species comparative studies to elucidate the exact mechanisms underlying these variations (Rauner et al. 2018, Hughes, in press). Moreover, the mammary gland field will greatly benefit from a comparative approach to mammary cancer using non-traditional model organisms, based on the observation that some mammals, such as horses, have a remarkably low incidence of mammary cancer (Bussche et al. 2017, Boyce and Goodwin 2017), despite their mammary gland closely resembling the human breast. Both species have mammary parenchyma surrounded by dense fibrous stroma, produce milk with a similar protein and fat content, and have terminal duct lobular units (TLDUs) as opposed to the terminal end buds (TEBs) in mice (Akers 2002, Spaas et al. 2012, Sharifi-Rad et al. 2013, Rauner et al. 2018). Collectively, MaOs from non-traditional model organisms for comparative research will provide further evolutionary insights into the development of the mammary gland across taxa and will uncover mechanisms underlying the variation in lactation strategies and mammary cancer incidence observed between mammals.

In this paper, we established and characterized MaOs from several non-traditional mammalian model organisms by using primary explants, in the form of mammary tissue fragments, that were embedded into a 3D matrix. To this end, we modified a method previously used to establish mouse MaOs (Ngyuen-Ngoc et al. 2015), to allow for the establishment of MaOs from cryopreserved mammary parenchyma embedded within a fibrous stroma. We first focused on establishing and characterizing equine mammary organoids (EqMaOs), and then expanded this protocol to additional mammals. Our salient findings were that EqMaOs can be successfully grown from tissue fragments, and that both budding and stage of non-budding EqMaOs, but not EqMaO diameter, is positively influenced by an increasing concentration of epidermal growth factor (EGF), which is required for branching morphogenesis in mouse MaOs and *in vivo* (Coleman et al. 1988, Sebastian et al. 1998, Simian et al. 2001).

Together, this study represents the first report on the establishment and characterization of MaOs from the horse and shows that this method is widely appliable to other non-traditional model organisms including cat, pig, deer, rabbit, and prairie vole.

## RESULTS

### Equine mammary organoids (EqMaOs) can be successfully generated from mammary tissue fragments (MTFs)

Since our initial attempts to establish equine mammary organoids (EqMaOs) from mammospheres and mammosphere-derived epithelial cells (MDECs) resulted in stellate colonies incapable of recapitulating mammary tissue polarity (data not shown) and attempts outside of our group using unsorted cells from canine mammospheres resulted in mammary organoids (MaOs) appearing stellate and lacking complex branching structures as well (Cocola et al. 2017), we assessed whether we could grow EqMaOs from mammary tissue fragments (MTFs) instead, using a modified protocol for growing MaOs from mice (Ngyuen-Ngoc et al. 2015). A key difference between the latter and our protocol was that the equine mammary tissues were collected and pre-digested using collagenase type II and cryopreserved for long-term storage (Fig. 1A), whereas the study using murine MTFs used fresh tissues. When equine mammary tissue was ready for MaO generation, it was thawed and minced, and digested again with collagenase type II to generate MTFs (Fig. 1B). Since the murine protocol was designed to isolate MTFs from adipose-rich mouse mammary fat pads (Ngyuen-Ngoc et al. 2015), we included some key modifications to our protocol to address the large amount of fibrous stroma in equine mammary tissues. First, tissues were strained using a metal sieve to separate small MTFs from larger pieces of undigested stroma immediately after the second collagenase digestion. Second, any trituration was avoided as this resulted in the release of fibrous stroma strings that contaminated the cultures and led to a poor MaO yield by breaking down MTFs into single cells. Next, we used a DNase treatment step to unstick the isolated MTFs from any contamination, followed by periodic centrifugation to dispose of these contaminants in the supernatant, keeping the fragments in a pellet. At this point, a successful versus a failed MTF isolation could be easily assessed by inspecting a small droplet of MTFs. After a successful MTF isolation, several large MTFs were readily observed (Fig. S1A). Common contaminants observed included single cells, nerve tissue fragments, and an unknown cellular contaminant (Fig. S1A). In contrast, a failed isolation would yield few and small MTFs with many single cells, likely as a result of over-mixing or triturating, as well as fibrous strings (Fig. S1B). MTFs from a successful isolation were then embedded in growth-factor reduced (GFR) Matrigel and supplemented with media. EqMaOs were cultured for 3 days and fixed in 4% PFA (Fig. 1B) and imaged using brightfield imaging for further analyses (Fig. 1C).

**Fig. 1.**
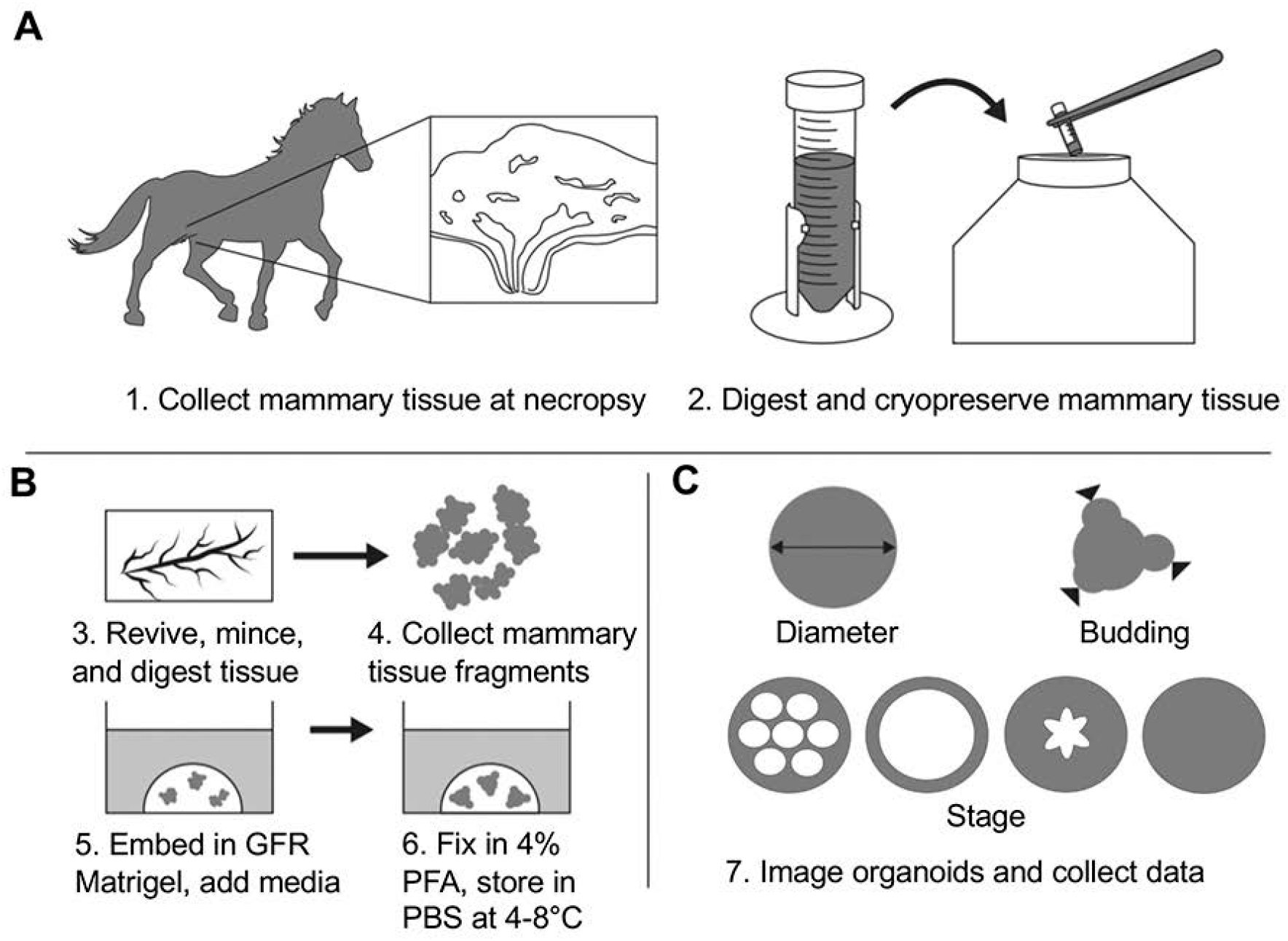
Graphical overview of tissue collection and cryopreservation, mammary tissue fragment isolation, mammary organoid (MaO) culturing and fixation, and analysis. **(A)** Mammary parenchyma was collected at necropsy, digested with enzyme solution A, and cryopreserved. **(B)** Mammary tissue was then thawed, minced using scissors, and digested again with enzyme solution B, which resulted in the isolation of mammary tissue fragments (MTFs). MTFs were embedded into growth factor reduced (GFR) Matrigel, supplemented with organoid media, and grown for 3 days. MaOs were fixed using paraformaldehyde (PFA) and stored at 4-8°C. **(C)** MaO data were collected using brightfield imaging and include MaO diameter, budding, and growth stage of non-budding MaOs.

An initial preliminary experiment using MTFs from Horse A (Table S1), showed that EqMaOs formed in both diluted (i.e. 3mg/ml, the lowest dilution needed to form a stiff 3D gel) and undiluted Matrigel (i.e. 9.4mg/ml), irrespective of the presence or absence of the growth factors epidermal growth factor (EGF) and fibroblast growth factor 2 (FGF2), alone or in combination (Fig. S2). However, EqMaOs in the diluted Matrigel conditions appeared to lose tissue polarity and cells began dissociating, resulting in a loss of structure (Fig. S2), and thus, we decided to grow EqMaOs in undiluted Matrigel.

### Increasing concentrations of epidermal growth factor (EGF) affect percentage of budding and stage, but not diameter, of equine mammary organoids (EqMaOs)

Our initial experiment showed that EqMaOs can form both in the presence or absence of growth factors, but since we noticed much larger EqMaOs in wells containing EGF (Fig. S2) we decided to evaluate the effects of increasing concentration of this growth factor on various parameters, such as MaO diameter, number of budding MaOs, and growth stage of non-budding MaOs, of EqMaOs established from three different mares. To this end, EqMaOs were cultured with increasing concentrations of EGF (2.5, 5, 10, and 20 nM) and a no treatment control and vehicle control, consisting of 0 nM EGF supplemented with plain medium or an equal volume of the vehicle (PBS with 10% 10mM acetic acid), respectively, were included as well. The latter was added to evaluate whether the vehicle that carries EGF has any effect on EqMaOs. After 3 days of culture, and fixation in 4% PFA, pictures were taken using brightfield imaging and collected data were further analyzed. One of the first things we noticed was that some larger EqMaOs, regardless of EGF concentration, appeared to be slow-growing and did not form buds when compared to the smaller, more typical EqMaOs. Since these large EqMaOs more closely resembled a MTF than an EqMaO, these rare, large EqMaOs were excluded from our analyses (Fig. 2A).

**Fig. 2.**
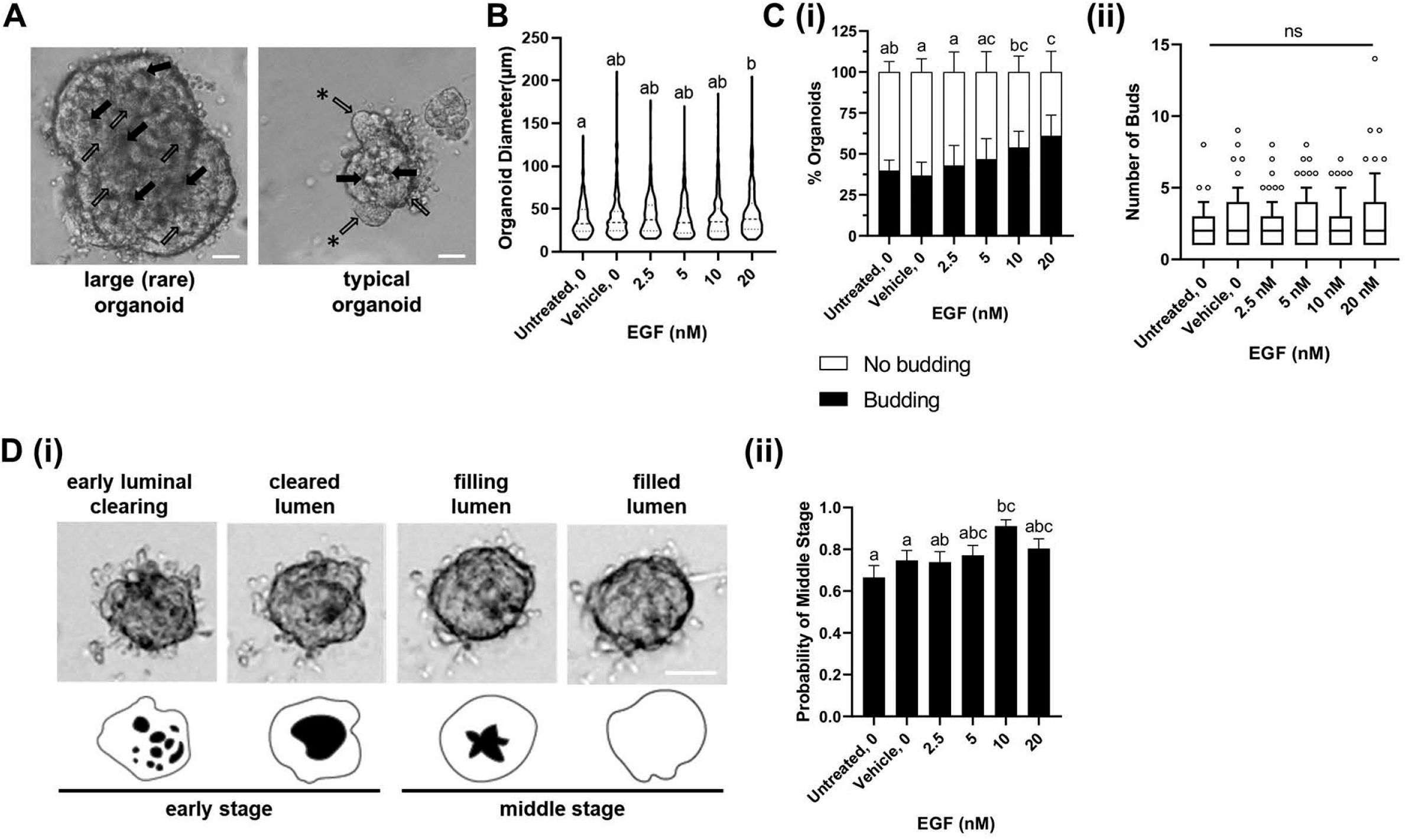
Equine mammary organoid (EqMaO) diameter, number of budding EqMaOs, and growth stage of non-budding EqMaOs, cultured in increasing concentration of epidermal growth factor (EGF). **(A)** Representative images of large (rare) and typical EqMaOs at 3 days of culture. Multi-layered epithelia (black arrows) surrounding luminal spaces (clear arrows). Asterisks indicate budding from the epithelia. **(B)** EqMaO diameter depicted by truncated violin plots (median ± 95% CI). Data were log-transformed and fit to a linear mixed-effects model, followed with a pairwise Tukey’s test. n=1390 EqMaOs. **(C)** Percentage of EqMaOs with 1 or more buds (budding) compared to EqMaOs with no buds (no budding). Data were fit to a binomial generalized linear mixed model, followed with a pairwise Tukey’s test. n=1390 EqMaOs. Error bars represent the standard deviation of the average percentage of budding and non-budding EqMaOs **(i)**. Number of buds per budding EqMaO (median ± 10-90^th^ percentile). Data were fit to a negative binomial generalized linear mixed model, followed with a pairwise Tukey’s test. n=621 EqMaOs. **(ii)**. **(D)** Sequential stages of growth in EqMaOs, represented by brightfield images (top) and cartoon schematics highlighting morphological changes in luminal space where early luminal clearing have a “honey-comb” appearance, the cleared lumen stage has a large circular lumen, a filling lumen has a characteristic “starfish” shape, and lastly a filled lumen will not have a lumen present (bottom). Scale bar represents 200 μm **(i)**. The probability of encountering “middle” stage versus “early” stage EqMaOs within non-budding EqMaOs. For data analysis, “early” and “middle” stages EqMaOs were fit to a binomial generalized mixed model, followed with a pairwise Tukey’s test. n= 769 EqMaOs. Error bars represent the standard error of the probability **(ii)**. For all data panels, different letters indicate significant differences (p < 0.05), ns=not significant, n=3 horses.

First, EqMaO diameter was evaluated and consistently showed a heavily right-skewed distribution with the majority of EqMaOs being 40 μm in diameter (Fig. 2B). No significant differences were found in diameter of EqMaOs cultured in wells with different concentrations of EGF, except a slight significant difference (p=0.0434) between the untreated control (0 nM) and 20 nM treatment (Fig. 2B). Next, we evaluated the percentage of budding EqMaOs and found that the culture wells without EGF, i.e. the untreated and vehicle control wells, contained approximately 40% EqMaOs with buds, with no statistically significant differences in percentage of budding EqMaOs between the untreated and vehicle control (Fig. 2C(i)). A significant difference in percentage of budding EqMaOs was observed between wells with 0 or 2.5 nM EGF, but not 5 or 10 nM EGF, and wells with 20 nM EGF (Fig. 2C(i)). Overall, there was an increase in percentage of budding EqMaOs when EGF concentration was increased. A more detailed analysis, where the number of buds per budding EqMaO were analyzed, did not show any statistically significant differences between the wells with varying concentration of EGF (Fig. 2C(ii)).

In parallel, we evaluated the growth stage of non-budding EqMaOs in more detail, based on data from murine MaOs showing that MaOs undergo different stages of tissue polarization and subsequent budding [21]. This process of sequential MaO growth, which we dub here as MaO “stage” for simplicity, begins with a multi-layered epithelium that undergoes luminal clearing, followed by progressive luminal filling, and lastly, the formation of buds where the lumen begins to form again resulting in an *in vivo*-like mammary structure [21]. Using time lapse imaging, we first verified that this type of MaO tissue polarization also occurred in EqMaOs (Movie S1). We divided the number of non-budding EqMaOs into four different stages: early luminal clearing, cleared lumen, filling lumen, and filled lumen; and then grouped them into two stages for statistical analysis, including early (consisting of early luminal clearing and cleared lumen) and middle (luminal filling and filled lumen) stages (Fig. 2D(i)). When comparing the number of middle stage EqMaOs to early stage EqMaOs across growth factor conditions and controls, we found that both 5 and 10 nM EGF were significantly different compared to both controls in producing middle stage EqMaOs, but we found no significant difference between 5, 10, and 20 nM EGF concentrations (Figure 2D(ii)).

### Time lapse imaging of equine mammary organoids (EqMaOs) reveals protrusions involved in organoid sensing and fusion

Irrespective of EGF concentration, we observed thin, finger-like protrusions, which are defined as putative actin-based cellular protrusions that are different from MaO buds, on EqMaOs from all 3 horses (Fig. 3A). Protrusions are thought to be involved in various processes such as cell sensation, cell-cell adhesion, and migration (Caswell and Zech, 2018). We hypothesized that these protrusions could be involved in MaO aggregation based on our observation that MaOs appeared to form protrusions in the direction of other MaOs which moved closer together during the 3-day culture period (Fig. 3B(i)), and in some cases, two MaOs completely fused together (Fig 3B(ii)).

**Fig. 3.**
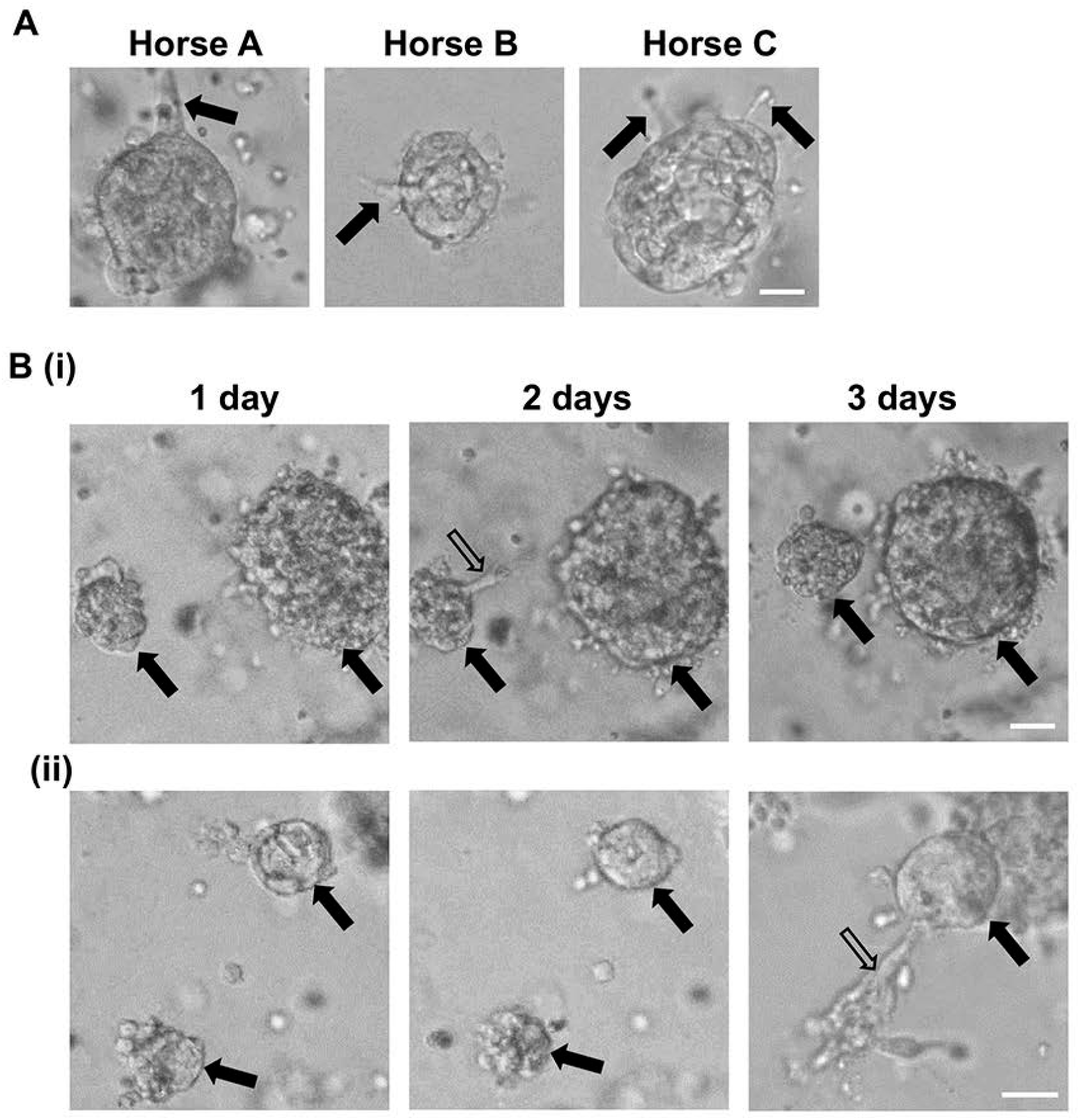
Equine mammary organoids (EqMaOs) have protrusions potentially involved in sensing and organoid fusion. **(A)** Representative images of a. finger-like protrusions on EqMaOs from three individual horses. **(B)** EqMaOs (black arrows) display protrusions (clear arrow) that grow towards other EqMaOs (i), and sometimes two individual EqMaOs (black arrows) fuse into one EqMaO through a finger-like protrusion (clear arrow) **(ii)**. Scale bars represent 50 μm.

To further investigate the activity of these protrusions, we used brightfield time lapse imaging to follow EqMaOs from Horse A in real time. We noticed that these protrusions were notably thinner and elongated when compared to buds (Fig. S3A), and they did not appear to grow from buds, suggesting that these protrusions are not involved in bud initiation/elongation, as has been previously demonstrated for murine MaOs (Ewald et al 2008). These finger-like protrusions in our cultures, however, did appear to have a sensory function by randomly probing the environment in various directions away from the EqMaO (Movie S2). We observed that when a finger-like protrusion from one EqMaO met another EqMaO (Fig. S3B(i)), a larger sheet-like protrusion would form between these two EqMaOs (Fig. S3B(ii)), followed by their fusion (Fig. S3B(iii)). The morphology of these protrusions appeared to be very dynamic (Movie S3), with sheet-like protrusions (Fig. S3C(i)) becoming finger-like protrusions (Fig. S3C(ii)), detaching from the EqMaO as a single cell (Fig. S3C(iii)), reattaching to the original EqMaO (Fig. S3C(iv)), and reforming back into a finger-like protrusion (Fig. S3C(v)). This suggests that EqMaO protrusions could be involved in functions beyond sensing, such as (i) adhesion between EqMaOs when the protrusions are sheet-like and/or cellular migration when the protrusions disseminate into single cells and (ii) in capturing escaped cells and reintegrating them back into the MaO, as has been reported previously for murine MaOs (Sirka et al. 2018).

### Equine mammary organoids (EqMaOs) are positive for markers of myoepithelial and luminal cells, and express β-lactoglobulin

We noticed that EqMaOs started to form 2D adherent monolayers after sinking through the Matrigel to the bottom of the culture plates, and interestingly, we could observe two distinct cell phenotypes consisting of cells with an elongated fibroblast-like and cells with a cobblestone morphology (Fig. S4), suggestive of the presence of myoepithelial and luminal cells, respectively. To confirm the nature of these two cell types, we used the myoepithelial marker CK-14 and the luminal marker CK-18 for immunohistochemistry (IHC) of EqMaOs from Horse A that were cultured in regular organoid media supplemented with 5 nM of EGF for 3 days. Since formalin-fixed paraffin-embedded (FFPE) equine mammary gland tissue was not available from Horse A, FFPE tissue from an age-matched, non-lactating mare, Horse D, was included as a positive control. As expected, CK14-positive signal was observed in basal cells within bilayered mammary epithelia and CK18-positive signal in the inner luminal cells (Fig. 4). EqMaOs showed a positive staining for both CK14 and CK18, although the positive signals were dispersed throughout the EqMaO and not localized to a defined compartment (Fig. 4). Next, we cultured EqMaOs in organoid media for an additional 3 days to obtain 6-day old EqMaOs, to evaluate whether further maturation of these EqMaOs could be obtained with a longer culture period. We observed that the positive CK14 signal became more compartmentalized into basal cells when compared to 3-day old EqMaOs; however, we still observed that CK18 was dispersed throughout the entire EqMaO (Fig. 4). Lastly, we stimulated 3-day old EqMaOs, cultured in regular organoid media supplemented with 5 nM of EGF, as described above, by replacing the regular organoid media with media supplemented with prolactin and devoid of EGF, and culturing for another 3 days. This approach is based on a previously established protocol (Sumbal et al. 2020), and we then stained these prolactin-stimulated, 6-day old EqMaOs with a marker for β-lactoglobulin, which is an abundant whey protein in equine milk (Sharifi-Rad et al. 2013, Wodas et al. 2020). We observed clear β-lactoglobulin-positive signal in EqMaOs irrespective of culture condition or culture period, but an additional positive stain was seen in the lumens of EqMaOs grown in the prolactin media, suggestive of milk secretion (Fig. 4). As expected, a weak β-lactoglobulin-positive signal was observed in the equine mammary gland tissue from the non-lactating mare (Fig. 4).

**Fig. 4.**
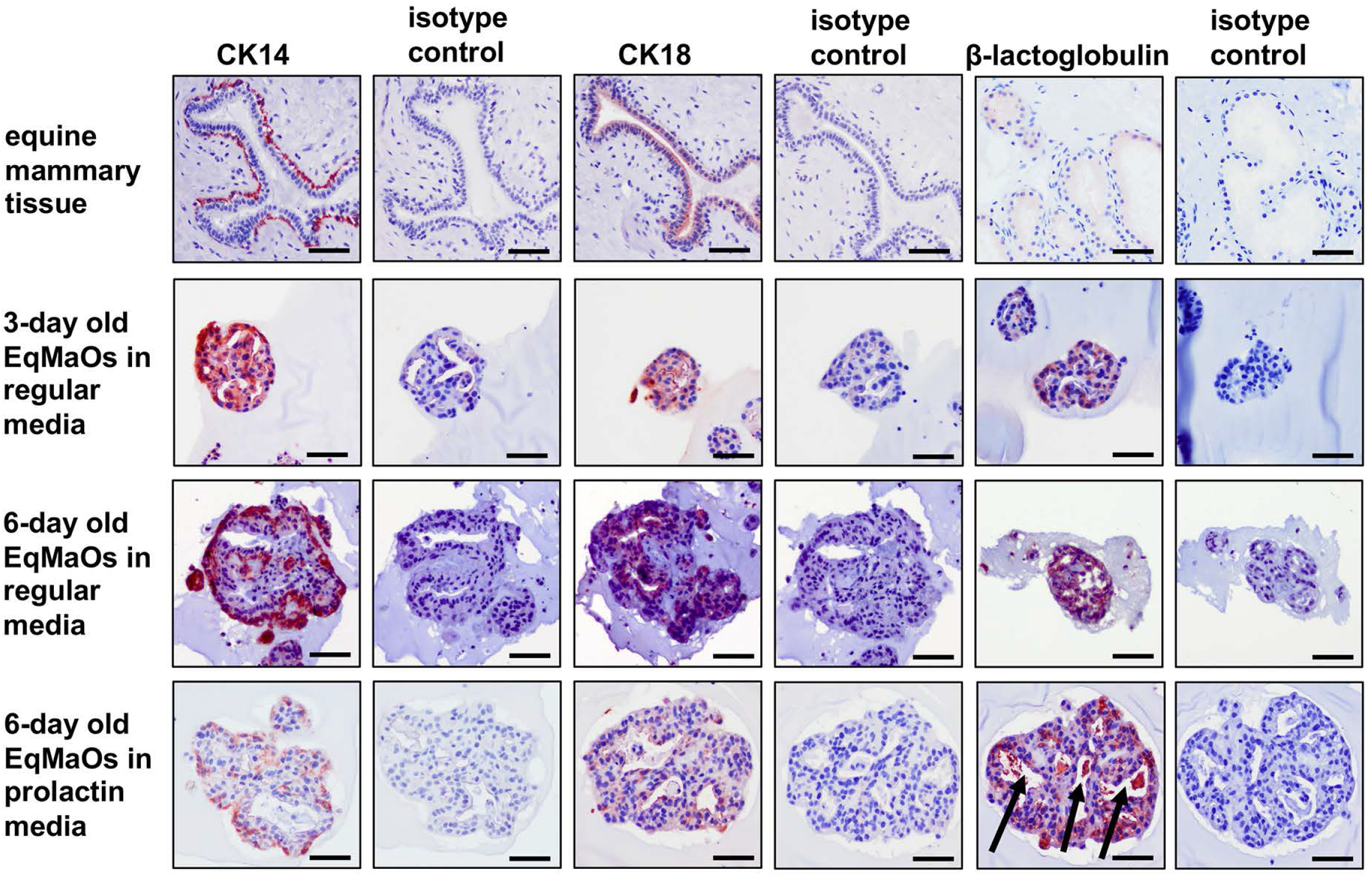
Immunohistochemistry (IHC) for cytokeratin (CK) 14, CK18, β-lactoglobulin. IHC was performed on equine mammary gland tissue from a non-lactating mare (top row), 3-day old (second row) or 6-day old (third row) equine mammary organoids (EqMaOs) grown in regular organoid media, and 6-day old EqMaOs grown in regular organoid media for 3 days, followed by an additional 3 days of culture in media supplemented with prolactin without EGF for (bottom row), using the myoepithelial maker CK14, the luminal marker CK18, and the whey protein β-lactoglobulin. Stainings with corresponding isotype controls were also performed. Arrows indicate proteinaceous material in EqMaO lumens. Scale bar represents 50 μm.

### Mammary organoids (MaOs) can be successfully established from other non-traditional model organisms using the mammary tissue fragment (MTF) approach

To evaluate whether our protocol could also be used to establish MaOs from other non-traditional model organisms, we attempted to culture MaOs from mammary gland tissues from cat (Felidae, FeMaO), pig (Suidae, SuMaO), deer (Cervidae, CeMaO), rabbit (Leporidae, LeMaO), and prairie vole (Cricetidae, CrMaO). MTFs were isolated from cryopreserved mammary gland tissue from these 5 mammals (Table S1) and all isolations contained MTFs, although the prairie vole and deer MTFs were rather sparse, despite multiple MTF isolation attempts (Fig. S5A). Similar to the horse, we observed different types of contamination in the MTFs isolations, including nerve tissue fragments and fibrous contamination across all species (Fig. S5B). In addition, muscle tissue fragments were noted as well, but only in the smaller mammals (e.g. cat, rabbit, and prairie vole) and not in the larger mammals (e.g. pig and deer) (Fig. S5B). When MTFs from these 5 mammals were cultured in undiluted Matrigel and organoid media supplemented with 5 nM EGF, MaOs were successfully established from all species and could be cultured for 6 days, except CrMaOs where most MaOs died off by day 3 of culture (Fig. 5A). FeMaOs, SuMaOs, and LeMaOs followed the same tissue morphogenesis patterns as seen with EqMaOs, consisting of a filled lumen by day 1, lumen formation and budding by days 3-4, and bud and MaO growth continuing into days 5-6 (Fig. 5A). MaO formation for these 3 species was, therefore, repeated with three biological replicates per species, which were fixed at day 3 of culture and analyzed for MaO diameter and percentage of budding MaOs. The MaO diameters of FeMaOs, SuMaOs, and LeMaOs, were all right skewed with average diameters of 65, 135, and 90 μm, respectively (Fig. 5B). The percentage of budding MaOs averaged around 30% and 31% for FeMaOs and LeMaOs, respectively, whereas SuMaOs had a much larger percentage of budding MaOs (71%) (Fig. 5C). CeMaOs survived up till 6 days in culture but remained small and did not go through the different morphogenesis stages, and CrMaOs had died by day 3 of culture (Fig. 5A), and were, therefore, not further analyzed quantitatively.

**Fig. 5.**
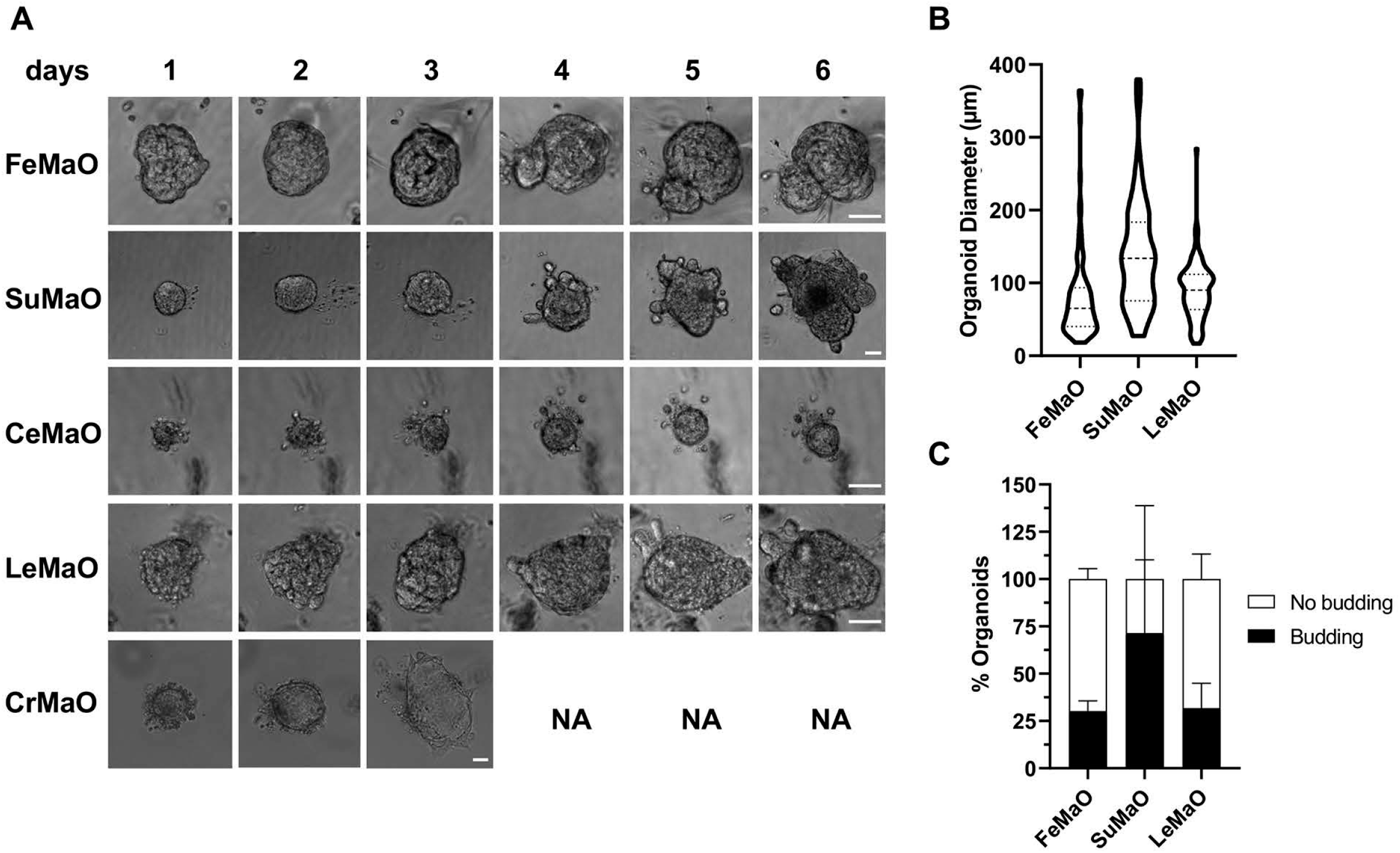
Mammary organoids (MaO) from additional non-traditional model organisms can be established using the mammary tissue fragment (MTF) methodology. **(A)** Representative images of cat (Felidae, FeMaO), pig (Suidae, SuMaO), deer (Cervidae, CeMaO), rabbit (Leporidae, LeMaO), and prairie vole (Cricetidae, CrMaO) MaOs cultured for 6 days. CrMaOs cultures were dead by 3 days of culture. Scale bars represent 50 μm. **(B)** MaO diameter depicted by truncated violin plots (median ± 95% CI). n=104, 44, and 99 MaOs for FeMaOs, SuMaOs, and LeMaOs, respectively. **(C)** Percentage of MaOs with 1 or more buds (budding) compared to MaOs with no buds (no budding). n=104, 44, and 99 MaOs for FeMaOs, SuMaOs, and LeMaOs, respectively. Error bars represent the standard deviation of the average percentage of budding and non-budding MaOs. n=3 per species.

To see if MaOs would survive passaging as well as to monitor further growth, we decided to passage 3-day old FeMaOs, SuMaOs, and LeMaOs, from one animal each by re-seeding them in fresh GFR Matrigel and culturing them for 6 days. In the original 3-day old cultures (passage 0), we observed both rounded, finger-like protrusions and spiny, mesenchymal-like protrusions on MaOs from all 3 mammals (Fig. S6A). In the passaged, 6-day old cultures (passage 1), several additional and interesting observations were made. First, we noticed that MaOs from FeMaOs and LeMaOs, but not SuMaOs, were able to develop secondary branches from a primary mammary bud (Fig. S6B). Second, we observed that MaOs from all 3 species formed alveolar-like structures, which were bubble-like outgrowths in FeMaOs or SuMaOs and more branch-like outgrowths in LeMaOs (Fig. S6B). When MaOs from all 3 species sank through the Matrigel to the bottom of the culture plates, they started to form 2D adherent monolayers containing cells with both fibroblast-like and cobblestone-like morphologies (Fig. S7), similar to what we observed previously with EqMaOs (Fig. S4) and thus, indicating that these MaOs also consist of myoepithelial and luminal cell types.

Collectively, the development of secondary branches and alveolar-like structures in passaged MaOs demonstrate that our MaO model is capable of recapitulating complex *in vivo*-like tubule-alveolar structures of the mammary gland.

## Discussion

There is an increasing recognition in the field of mammary gland biology and neoplasia to expand research into using understudied, non-traditional mammal species, as this allows for comparative studies resulting into newly gained knowledge based on the variation in mammary gland development and cancer incidence across mammals (Rauner et al. 2020, Harman et al. 2020, Hughes, 2020). Here, we report for the first time the establishment and characterization of equine (Eq), feline (Fe), porcine (Su), and leporine (Le) mammary organoids (MaOs) and propose these MaOs as useful models for comparative studies investigating species-level differences in mammary development, natural cancer variation, mammary gland anatomy and physiology, as well as mammary gland evolution.

Specifically, we describe a protocol using mammary tissue fragments (MTFs), synonymous to previously described murine “epithelial pieces” (Fata et al. 2007) and “primary mammary organoids” (Ngyuen-Ngoc et al. 2015) or “mammary organoids” in bovine (Martignani et al. 2018), which can be embedded in Matrigel and used to grow MaOs for an array of non-traditional model organisms, including horses. To avoid confusion in the description of mammary fragments as “organoids”, we propose that the word “organoid” should be restricted to 3D-matrix-embedded mini-organs and not be used for mammary tissue fragments. This method of using MTFs is a straightforward and accepted way to study mammary gland development, because the MaOs form quickly in culture and are able to form complex structures such as mammary ducts and alveolar structures (Nguyen-Ngoc et al. 2015). Other methods, such as embedding single cells from dissociated mammospheres in a 3D matrix to form MaOs have been used as well, but these MaOs appear to lack complex structures, or form stellate colonies (Cocola et al. 2015), an observation we made ourselves using equine mammospheres. In contrast, the use of sorted CD49f high and low cells to establish mammospheres from marmosets, which were then grown into MaOs, seemed more successful since no stellate colonies were observed. However, these marmoset MaOs did not appear to have extensive branching nor alveolar structures (Wu et al. 2016). The use of stem cells to generate MaOs is another approach (Rosenbluth et al. 2020), but this is only feasible for those species with established cell sorting protocols for adult mammary gland stem cells and/or reprogramming protocols for induced pluripotent stem cells (IPSCs). To date, sorting protocols for adult mammary gland stem cells are primarily established for mouse (Stingl et al. 2006), human (Gudjonsson et al. 2002, Qu et al. 2017), and cow (Cravero et al. 2015); and although IPSC technology has been applied to various mammals, both domesticated and wild (Stanton et al. 2019), it is not routinely used for generation of MaOs, at least not to our knowledge.

An in-depth analysis was performed on EqMaOs, due to our interest in the naturally low incidence of mammary cancer in horses, despite the equine mammary gland being similar to the human breast in relation to development, anatomy, and function (Akers 2002, Spaas et al. 2012, Sharifi-Rad et al. 2013, Rauner et al. 2018). Several morphological parameters, including diameter, budding (Simian et al. 2001), and stage of non-budding MaOs, were evaluated in 3-day old EqMaOs in the context of increasing concentrations of epidermal growth factor (EGF) (Coleman et al. 1988, Sebastian et al. 1998). This 3-day culture period was based on murine MaO studies, where most MaO budding was found to occur between 2-3 days (Fata et al. 2007). To our knowledge, we are the first to formally report on MaO diameter, and observed a wide, albeit heavily skewed, range of EqMaO diameters irrespective of increasing EGF concentrations. Interestingly, we consistently observed a few large EqMaOs (>100 μm) in our cultures as well, but those did not readily undergo tissue polarization to form MaOs leading us to hypothesize that increased size could limit nutrient diffusion or that larger MaOs have more cells and need more time to rearrange in culture. The need for optimization of organoid size is well-recognized in non-luminal organoid models, such as cerebral and liver organoids, where necrosis is observed due to decreased nutrient diffusion to the innermost cells (Akkerman 2017, Grebenyuk et al. 2019). In contrast to MaO diameter, we did notice that EGF supplementation increased the number of budding EqMaOs significantly, a finding similar to studies with murine MaOs (Simian et al. 2001, Fata et al. 2007).

We had variable success in growing MaOs from additional, non-traditional model organisms, with MaOs from domesticated animals such as cats, pigs, and rabbits, readily forming and being healthy, whereas MaOs established from wild animals such as deer and prairie voles not developing and maturing or quickly dying off in culture, respectively. The latter could be due to technical difficulties with dissecting mammary gland parenchyma and/or isolating sufficient MTFs from these species. Further analysis of FeMaOs, SuMaOs, and LeMaOs, in comparison to our previously established EqMaOs, showed that (i) MaOs from cat, pig, and rabbit, were on average larger compared to those from horse and (ii) SuMaOs had the highest percentage of budding MaOs from all species, although there appeared to be large variability between biological replicates.

The presence of abundant protrusions and disseminating cells has been previously reported on mouse MaOs grown in collagen, but not Matrigel (Ngyuen-Ngoc 2012). In contrast, we observed many protrusions and disseminating cells on our MaOs from non-traditional model organisms grown in Matrigel. Compared to Matrigel, collagen is known to induce “invasive” phenotypes in MaOs such as cell dissemination and protrusions (Ngyuen-Ngoc et al. 2012, Sumbal et al. 2020). Interestingly, when we tried to grow mouse MaOs using our protocol involving Matrigel, we did observe some protrusions, albeit not nearly as much as on the MaOs we established from the non-traditional model organisms (data not shown). The ability of MaOs from non-traditional species to readily form protrusions and disseminating cells in Matrigel may be due to evolutionary differences in the branching morphogenesis *in vivo* between animals with a fatty (rodents) or fibrous stroma (e.g. horses, cats, pigs, rabbits). It could be hypothesized that the inherent properties of mammary epithelium between mice and these non-traditional model organisms differ, such as their mechanisms for branching morphogenesis, leading to these observable differences in behavior across different 3D matrices. Indeed, a review from Sumbal and colleagues (2020) points out that human and mouse MaOs may have different means of growth through a 3D matrix, involving an “invasive” and “non-invasive” mechanism, respectively. The function of these protrusions and disseminating cells, and how they interact with the microenvironment, has been a recent area of interest, uncovering that normal branching morphogenesis mechanisms are partially co-opted in mammary cancer for invasion and that the myoepithelium serves as a barrier to cell dissemination (Sirka et al. 2018, Feinberg et al. 2018, Sumbal et al. 2020). Although we currently don’t know the exact nature of the protrusions we observed on the MaOs from these non-traditional model organisms, nor what their specific functions are, future research studying these protrusions in greater depth will provide additional insights into evolutionary and developmental biology.

Taken together, the establishment of MaOs from non-traditional model organisms, as described in this study, will provide a new resource for comparative and/or stand-alone mammary developmental-, evolutionary-, and cancer-focused studies across species.

## MATERIAL AND METHODS

### Ethics statement

Since all tissues were collected after euthanasia/culling, no IACUC approval is needed.

### Mammary tissue collection

Mammary tissues were collected from various mammals, euthanized/culled for reasons unrelated to this study, and obtained through research labs at Cornell University College of Veterinary Medicine (horse, pig, rabbit, and vole), local hunters (deer), or Marshall BioResources (cats; North Rose, NY). Details related to breed, age, and history, of these animals are provided in Supplementary Table 1. Mammary gland parenchyma was collected using sterile techniques and placed in PBS for a maximum of 1 to 2 h on ice until processing. Tissue was washed with Hank’s Balanced Salt solution (HBSS) (Thermo Fisher, Waltham, MA) + 5% antibiotic-antimycotic (Thermo Fisher). Tissue was minced using sterile scissors into 3-5 mm^3^ pieces and digested in Enzyme solution A, composed of 1.2 mg/mL collagenase type II (Worthington Biochemical Corporation, Lakewood, NJ), 100 U/mL hyaluronidase (Sigma Aldrich, St Louis, MO), 5% fetal bovine serum (FBS) (Atlanta Biological, Flowery Branch, GA), 5 μg/mL human recombinant insulin (Thermo Fisher), and 1 μg/mL hydrocortisone (Sigma Aldrich), in 1:1 DMEM/Ham’s F12 (Corning, Acton, MA), for 3 h rocking at 37°C. After incubation, tissues were softened by triturating with a wide-bore pipette and washed three times with 2% HBSS with 2% FBS (HF) with periodic centrifugation at 1,250 xg to remove the supernatant. Tissues were resuspended in freezing media (FBS + 10% sterile DMSO), aliquoted into 2 mL cryovials (Corning), and transferred to liquid nitrogen for long-term storage.

### Mammary tissue fragments (MTFs) isolation

Cryopreserved mammary tissues were thawed, washed with sterile PBS, and centrifuged to remove supernatants. Tissues were minced using sterile scissors into 1 mm^3^ pieces and digested in Enzyme solution B, composed of 2 mg/mL collagenase type II, 5% FBS, 5 μg/mL insulin, 50 μg/mL gentamycin (Thermo Fisher), and 2 mg/mL trypsin (Corning), in 1:1 DMEM/Ham’s F12, for 45-60 min rocking at 37°C. Beyond this point, all tubes and pipettes used to hold or transfer MTFs were coated with 0.1% bovine serum albumin (BSA) (Sigma Aldrich) solution in PBS to prevent MTFs sticking to surfaces. MTFs were collected using a metal sieve with around 500 μm size to strain out larger pieces of tissue and washed with 10 mL 1:1 DMEM/Ham’s F12. Next, 50-150 μl of 5 mg/mL DNase (Sigma Aldrich) was added and slowly hand rocked for 5 min, causing MTFs to detach from contaminating single cells and fibrous contamination, which were then further separated via 4 periodic centrifugation steps at 1,250 xg. After carefully removing the supernatant above the MTF pellet and pipetting out visible fibrous material during resuspension of pellet, MTFs were finally resuspended in fresh 1:1 DMEM/Ham’s F12. MTFs were counted, exactly as previously described (Nguyen-Ngoc 2015).

### Mammary organoid (MaO) culturing, definitions, and data collection

MTFs were counted and embedded into 5 μl of 9.4 mg/mL Growth Factor Reduced (GFR) Matrigel (Corning) at a density of 10 MTFs/μl in 5 μl domes in a 96-well plastic bottom plate (Corning) to form MaOs. MaOs were kept at 5% CO2, 37°C, at 80% humidity. MaOs were cultured for 3 days, unless where indicated otherwise, in organoid media composed of 1% penicillin-streptomycin (Thermo Fisher), 1% insulin-transferrin-selenium (Thermo Fisher) in 1:1 DMEM/Ham’s F12 and varying concentrations (i.e. 0, 2.5, 5, 10, or 20 nM) of epidermal growth factor (EGF) (Sigma Aldrich). Each of these groups were done in triplicate wells. Since EGF was diluted in a 10 mM acetic acid + 0.1% BSA solution in PBS, a vehicle control consisting of 10% 10 mM acetic acid + 0.1% BSA in PBS was included in all experiments. After the 3-day culture period, MaOs were fixed in 4% paraformaldehyde (PFA) (Sigma Aldrich) for 10 min, washed three times with PBS for 10 min, and stored at 4°C; or passaged into new GFR Matrigel and supplemented with fresh organoid media. For the latter, GFR Matrigel was mechanically broken down using a pipette tip and liquified in chilled 4°C DMEM/Ham’s F12, after which MaOs were re-embedded in an equal volume of fresh GFR Matrigel and cultured for 6 days and passed again. Brightfield images of MaOs were obtained using a Zeiss Axio Observer inverted microscope, an Olympus CKX41 inverted microscope fitted with a Lumenera Infinity 2-3C camera, or a ZOE Fluorescent Cell Imager. The Zeiss Axio Observer inverted microscope was used to collect images for statistical analyses and one randomly selected field within 3 individual wells of the same condition were used for data collection. Zeiss; ZEN 3.1 (blue edition) software was used to measure MaO diameter. Both the percentage of budding MaOs and the number of buds per MaO were quantified. Buds are the structures that give rise to mammary ducts and undergo elongation from the central spherical body of the MaO (Simian et al. 2001, Fata et al. 2007, Ewald et al. 2008). An MaO was considered budding if it had one or more buds. Classification of sequential stages in MaO growth, dubbed MaO “stage” in non-budding MaOs, was assessed first by the lack of budding and second by the appearance of the lumen. To assess the lumen, MaOs were categorized as followed: the early luminal clearing stage had a “honey-comb” appearance, the cleared lumen stage had a large circular lumen, the filling lumen stage had a characteristic “starfish” shape, and the filled lumen stage did not have a lumen present. MaO diameter, budding, and stage, were the dependent variables, while the increasing concentration of EGF was the independent variable. All data were collected by one, unblinded, observer. MaOs which still resembled mammary tissue fragments and had not entered the first stage of early luminal clearing were excluded from statistical analyses.

### Time-lapse Imaging

To provide a single plane for MaOs to rest on during time-lapse imaging, 200 μl of undiluted GFR Matrigel (9.4 mg/ml) was plated onto a pre-incubated (37°C) MatTek dish (MatTek Corporation, Ashland, MA) and incubated for 30 min at 37°C to polymerize. Cryopreserved MTFs from Horse A were thawed and revived, washed with 10 mL sterile PBS by centrifuging at 1,250 xg until pelleted, and supernatant was removed. The MTFs were mixed into 100 μl of diluted GFR Matrigel (3.0 mg/ml) and plated on top of the polymerized undiluted Matrigel. MaOs sunk in the diluted GFR Matrigel until they reached the undiluted GFR Matrigel, putting all MaOs within the same viewing plane for time-lapse imaging. Brightfield images were taken every 15 min at 5x magnification on a Zeiss Axio Observer inverted microscope fitted with a built-in incubator maintained at 37°C with 5% CO2.

### Immunohistochemistry (IHC)

Equine MaOs (EqMaOs) from Horse A were prepared by growing them in 50 μl GFR Matrigel at a density of 10 MTFs/μl in 24-well plastic bottom plates (Corning) for 3 days in regular organoid media supplemented with 5 nM EGF. EqMaOs were then cultured for another 3 days in either regular organoid media or in organoid media without EGF, supplemented with 1 ug/mL sheep recombinant prolactin. EqMaOs were fixed in 4% PFA for 20 min, washed 3x with PBS for 10 min, and stored at 4°C until further analysis. To prepare EqMaOs for paraffin embedding, HistoGel (Thermo Fisher) was liquified in a 65°C water bath, a 3D-printed hollow cylindrical mold was placed on top of the fixed Matrigel dome and liquid HistoGel was poured into the mold. Using a pipette tip, Matrigel and EqMaOs were resuspended in the HistoGel by gently scraping them from the glass slide they were cultured on, and then placed on ice to solidify. The HistoGel-containing EqMaOs were then dehydrated by being placed in 30%, 50%, and 70% ethanol in distilled water for 15 min, to remove the HistoGel, and were then stored in 70% ethanol. The EqMaOs were sent to the Cornell University Histology Core for processing. Paraffin embedded EqMaO sections were deparaffinized and rehydrated. EqMaOs were permeabilized with Tris buffered saline (TBS) + 0.5% Tween-20 (Sigma Aldrich) for 10 min at room temperature (RT) and washed with wash buffer (TBS + 0.1% Tween-20) 3 times for 5 min. Antigen retrieval was performed using sodium citrate buffer, pH=6, by microwaving on low power for 20 min, followed by 3 washes for 5 min with wash buffer. EqMaOs were blocked with anti-goat blocking solution (TBS + 1% BSA +1-% normal goat serum (Thermo Fisher). Anti-cytokeratin (CK) 14 anti-CK 18 antibodies (Abcam, ab7800 and ab668, respectively), a myoepithelial and luminal marker, respectively, were diluted 1:100 in diluent solution (TBS + 1% BSA), and anti-β-lactoglobulin (Abcam, ab112893) antibodies were diluted 1:1000. Equal concentrations of either mouse IgG (Abcam, ab18443) (CK14 and CK18) or rabbit IgG (Abcam, ab172730) (β-lactoglobulin) were used as isotype controls. Primary antibody incubations were carried out at 4°C overnight, followed by 3 5-min washes with wash buffer. A 0.3% hydrogen peroxide solution in TBS was applied for 15 min and slides were washed once for 5 min with wash buffer. Secondary antibodies, either goat anti-mouse IgG-HRP (Jackson, 115-035-062) or goat anti-rabbit IgG-HRP (Jackson, 111-035-144), respectively, were diluted 1:500 and incubated for 1 h at RT, followed by 3 5-min washes with wash buffer. Where applicable, biotin-labeled anti-mouse IgG antibodies (Jackson) were used, diluted 1:500 and incubated for 1 h at RT, followed by incubation with streptavidin-HRP, diluted to 2 μg/ml, for 1 h at RT, to increase signal. An AEC kit (Thermo Fisher) was used as a chromogen and slides were counterstained with Gills 2 hematoxylin (Thermo Fisher). Since mammary tissue sections from Horse A were unavailable, paraffin-embedded mammary tissue sections from an age-matched, non-lactating mare (Horse D) were included as positive control.

### Statistical Analysis

Statistical analyses were carried out in RStudio (v 1.1.456). For analysis of EqMaO morphological parameters, including diameter, budding, and stage, we accounted for random effects between individual horses (n=3, Supplementary Table 1). Pairwise comparisons were done using Tukey’s method. Assumptions including normality of the residuals and constant variance were checked. An alpha level of 0.05 was used for all tests. For budding analysis, we grouped data into “budding” or “not budding” categories and used a binomial generalized linear mixed model (GLMM). We then performed a follow-up analysis on the budding EqMaOs to see if the was a difference in bud count per condition. To this end, because residuals were right-skewed, we fit the data to a negative binomial GLMM. To analyze differences in EqMaO stage within the non-budding EqMaOs, we grouped data into either “early” or “middle” stage EqMaOs and used a binomial GLMM to compare the probability of encountering middle stage EqMaOs versus early stage EqMaOs. Similar analyses were carried out for MaOs from other species, where applicable.

## Acknowledgements

We are very grateful to Sarah Johnston for the graphic design of Figure 1 and Stephen Parry for his assistance with statistical analysis. We thank Dr. Rebecca Harman and James Miller for help with immunohistochemistry. We also like to thank Dr. Chinatsu Mukai for her assistance in the time-lapse imaging of mammary organoids.

## SUPPLEMENTARY MOVIE LEGENDS

**Movie S1.** Brightfield time-lapse movie of an equine mammary tissue fragment (MTF) cultured in growth factor reduced (GFR) Matrigel with organoid media supplemented with 5 nM epidermal growth factor (EGF) that undergoes tissue morphogenesis into an equine mammary organoid (EqMaO) with budding mammary ducts.

**Movie S2.** Brightfield time-lapse movie of two equine mammary organoids (EqMaOs) showing finger-like protrusions that probe the environment and appear to be involved in organoid fusion.

**Movie S3.** Brightfield time-lapse movie showing the dynamic morphology and behavior of protrusions on equine mammary organoids (EqMaOs).

**Supplementary Table 1.** Mammary tissue samples from female mammals used in this study.

| **Name** | **Breed** | **Age** | **History** |
| --- | --- | --- | --- |
| Horse A | Saddlebred | 7 years | Unknown. Mare suspected of previous laccation. Mammary gland was involuted at time of tissue collection. |
| Horse B | Warmblood | 4 years | No pregnancies, intact. Surgical procedure with general anesthesia to induce cartilage lesion in stifle. |
| Horse C | Thoroughbred | 3 years | No pregnancies, intact. Surgical procedure with general anesthesia to induce cartilage lesion in stifle. |
| Horse D | Warmblood | 4 years | No pregnancies, intact. Surgical procedure with general anesthesia to induce cartilage lesion in stifle. |
| Cat A | Unknown | < 3 years | Pathogen-free colony-bred for research purposes. |
| Cat B | Unknown | < 3 years | Pathogen-free colony-bred for research purposes. |
| Cat C | Unknown | < 3 years | Pathogen-free colony-bred for research purposes. |
| Pig A | Mixed breed, predominantly Hampshire | 5 months | Virgin pig used for study on joints. Surgical procedure under general anesthesia. |
| Pig B | Mixed breed, predominantly Hampshire | 5 months | Virgin pig used for study on joints. Surgical procedure under general anesthesia. |
| Pig C | Mixed breed, predominantly Hampshire | 4 months | Virgin pig used for research purposes.. |
| Rabbit A | New Zealand White | 16 weeks | Intact virgins used for a physiology class. |
| Rabbit B | New Zealand White | 16 weeks | Intact virgins used for a physiology class. |
| Rabbit C | New Zealand White | 16 weeks | Intact virgins used for a physiology class. |
| Deer | White-tailed deer | Unknown | Adult female, likely parous but not suspected of lactating at time of tissue collection. |
| Prairie vole | NA | Unknown | Pregant at time of tissue collection. Two animals were pooled into one sample. |

**Fig. S1.**
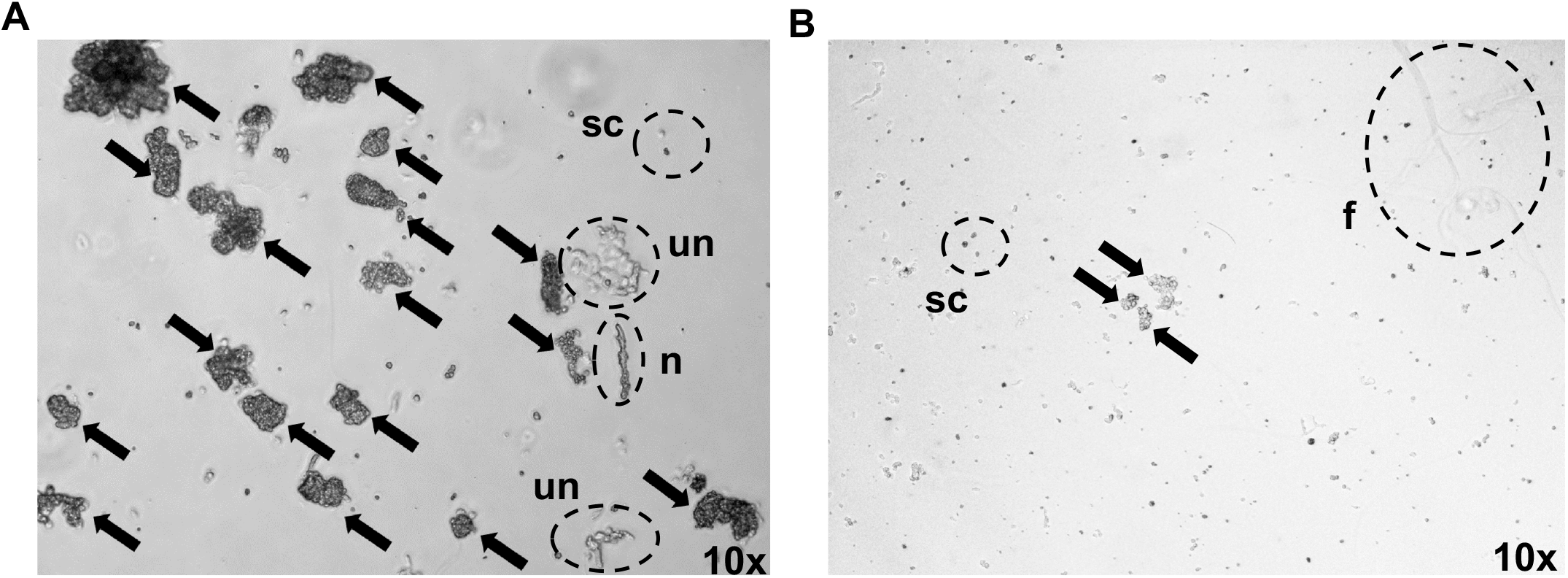
**(A)** Representative image (10X) of a successful equine mammary tissue fragment (MTF) isolation showing large and many MTFs (black arrows), in addition to commonly observed contaminants including single cells (sc), nerve tissue fragments (n), and unknown fibrous/cellular clusters (un). **(B)** Representative image (10X) of a failed equine MTF isolation showing small and few MTFs (black arrows), as well as many single cells (sc) and fibrous contamination (f).

**Fig. S2.**
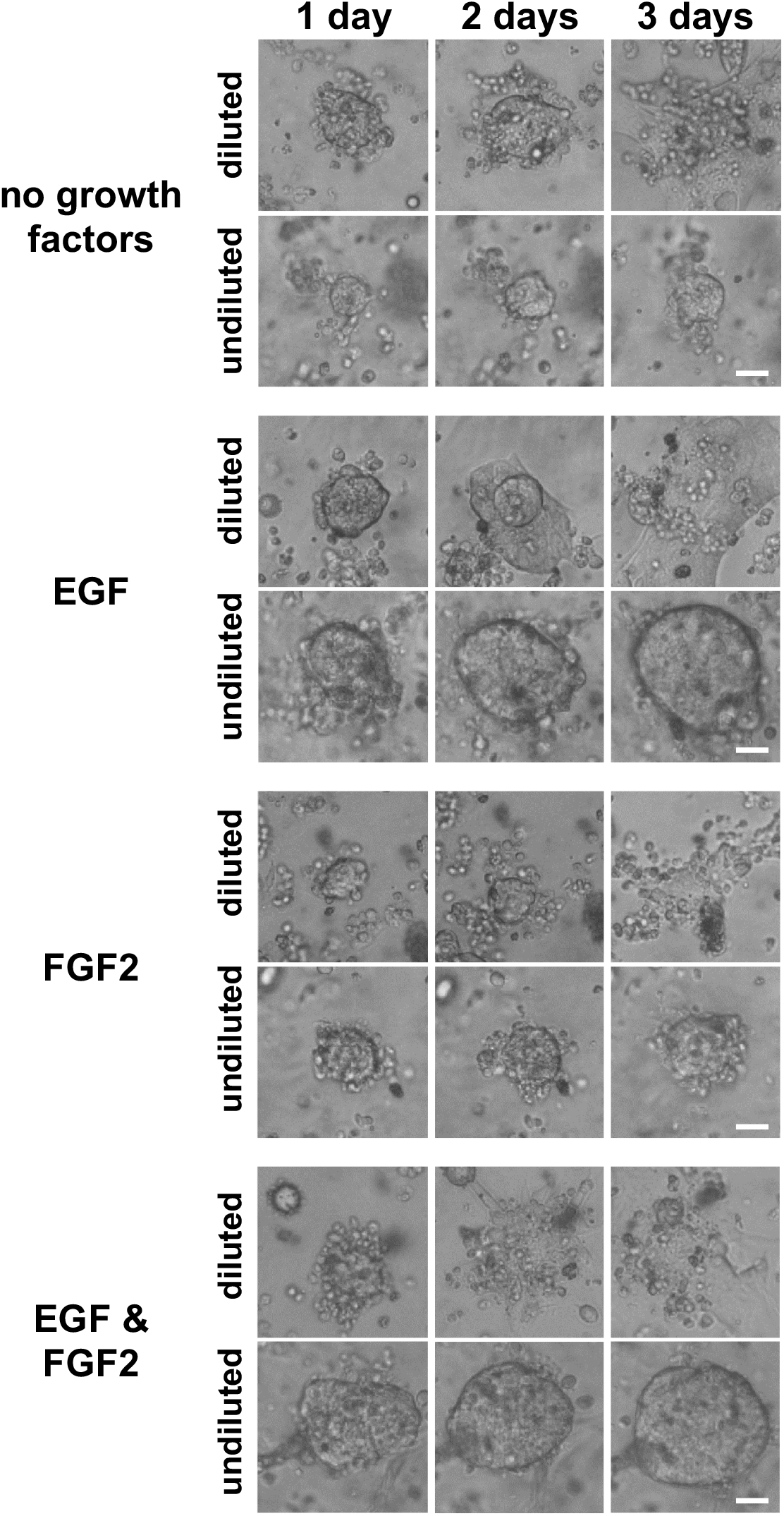
Equine mammary organoids (EqMaOs) grown for 3 days in either diluted growth factor reduced (GFR) Matrigel or undiluted GFR Matrigel and supplemented with organoid media containing 2.5 nM EGF; 2.5 nM FGF2; 2.5 nM EGF and 2.5 nM FGF2; or no growth factors. Scale bars represent 25 μm.

**Fig. S3.**
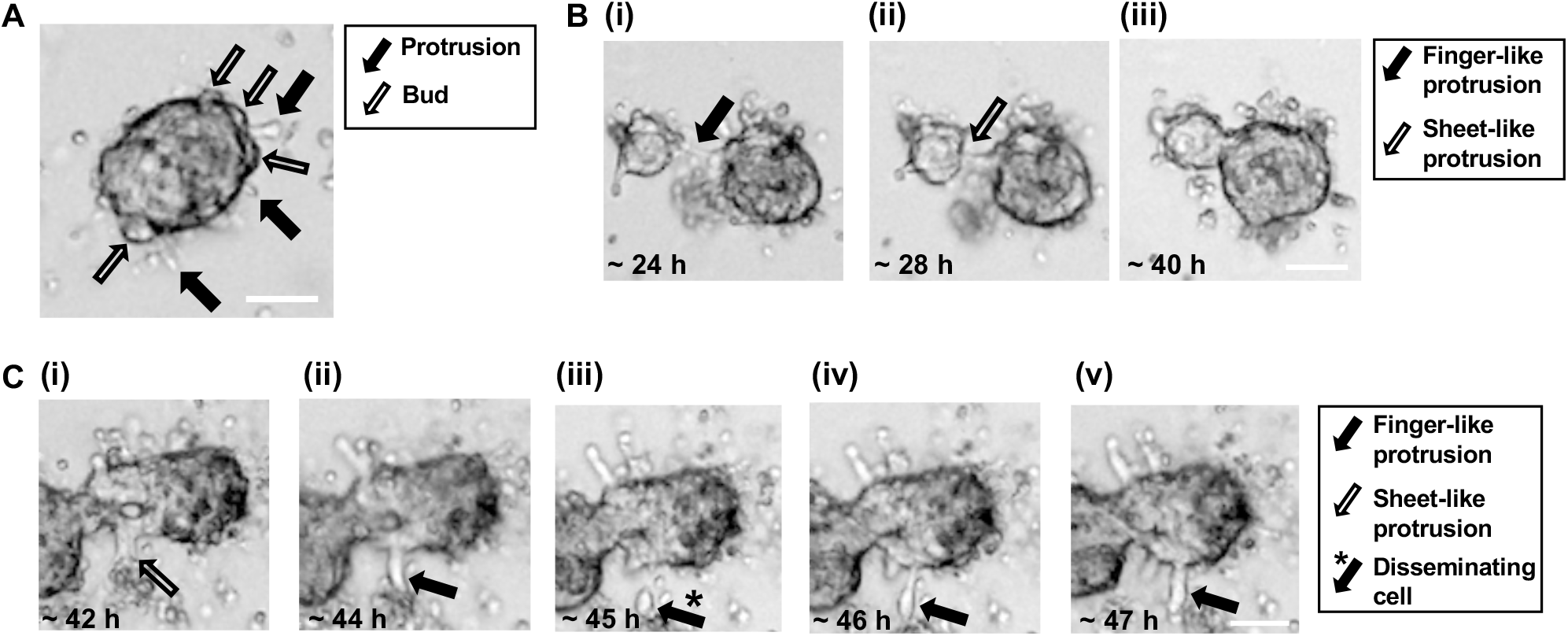
Protrusion morphology and function in EqMaOs. **(A)** Representative image of an EqMaO with both thin, elongated protrusions (black arrows) and thicker, shorter buds (clear arrows). **(B)** Representative images over time of two EqMaOs connected via a finger-like protrusion (black arrow) (~24 h, **i**), which then became thicker and sheet-like (clear arrow) creating a stronger bridge between the two organoids (~28 h, **ii**), to finally result in fusion of the two organoids (~40 h, **iii**). **(C)** Representative images of protrusion dynamics on EqMaOs. Sheet-like protrusions (clear arrow) around 42 h of culturing **(i),** became finger-like protrusions (black arrows) at 44 h **(ii)**, followed by dissemination into single cells (black arrow with *) around 42 h of culturing **(iii)**. These single cells reattached to the original organoid (black arrow) at 46 h **(iv)** to reform a finger-like protrusion (black arrow) at 47 h **(v)**. Scale bar represents 200 μm.

**Fig. S4.**
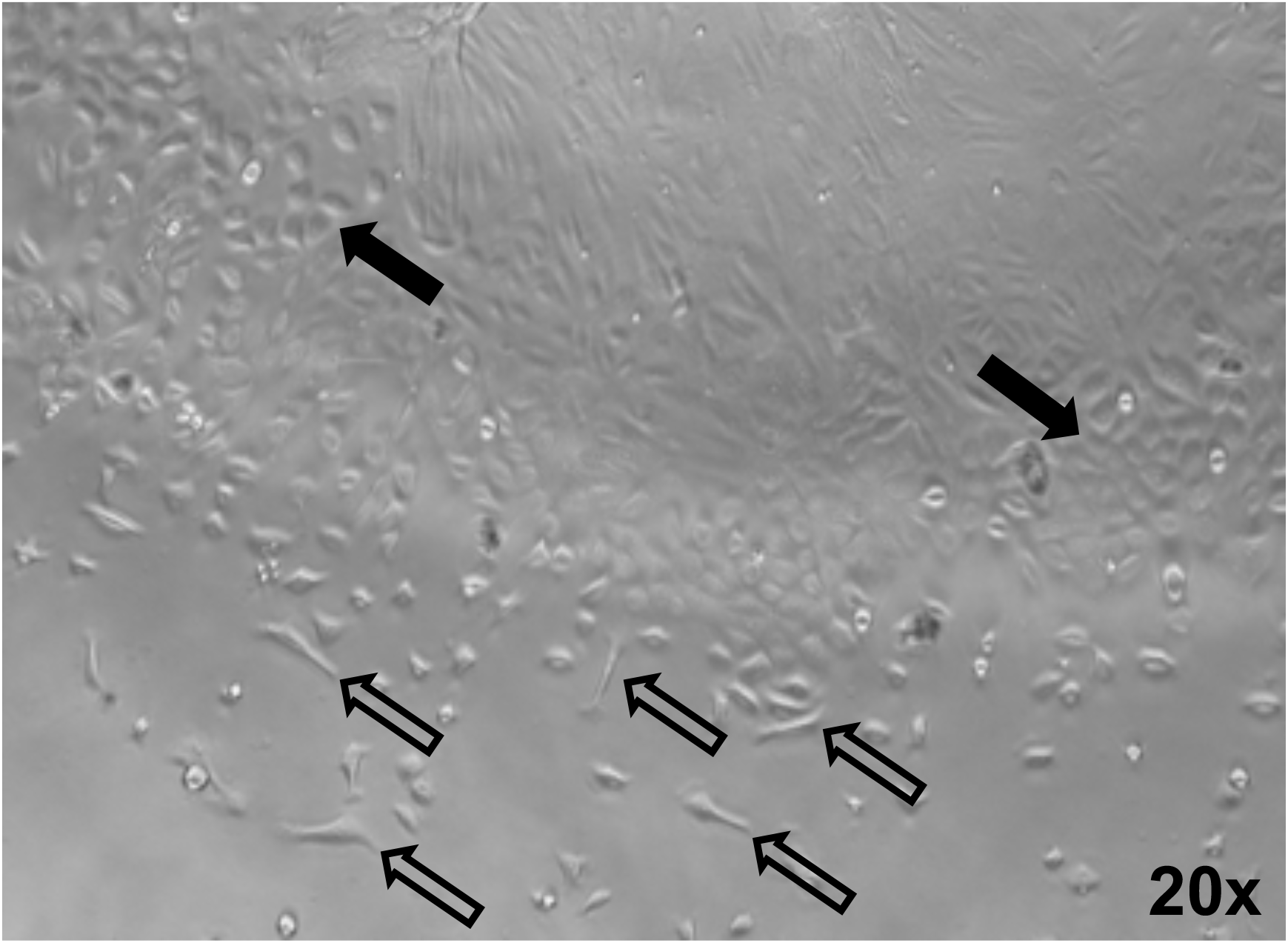
Representative image (20X) of equine adherent cells with both mesenchymal-like morphology (clear arrows) and cobblestone morphology (black arrows) after EqMaOs sank through the Matrigel to the bottom of the cell culture plates.

**Fig. S5.**
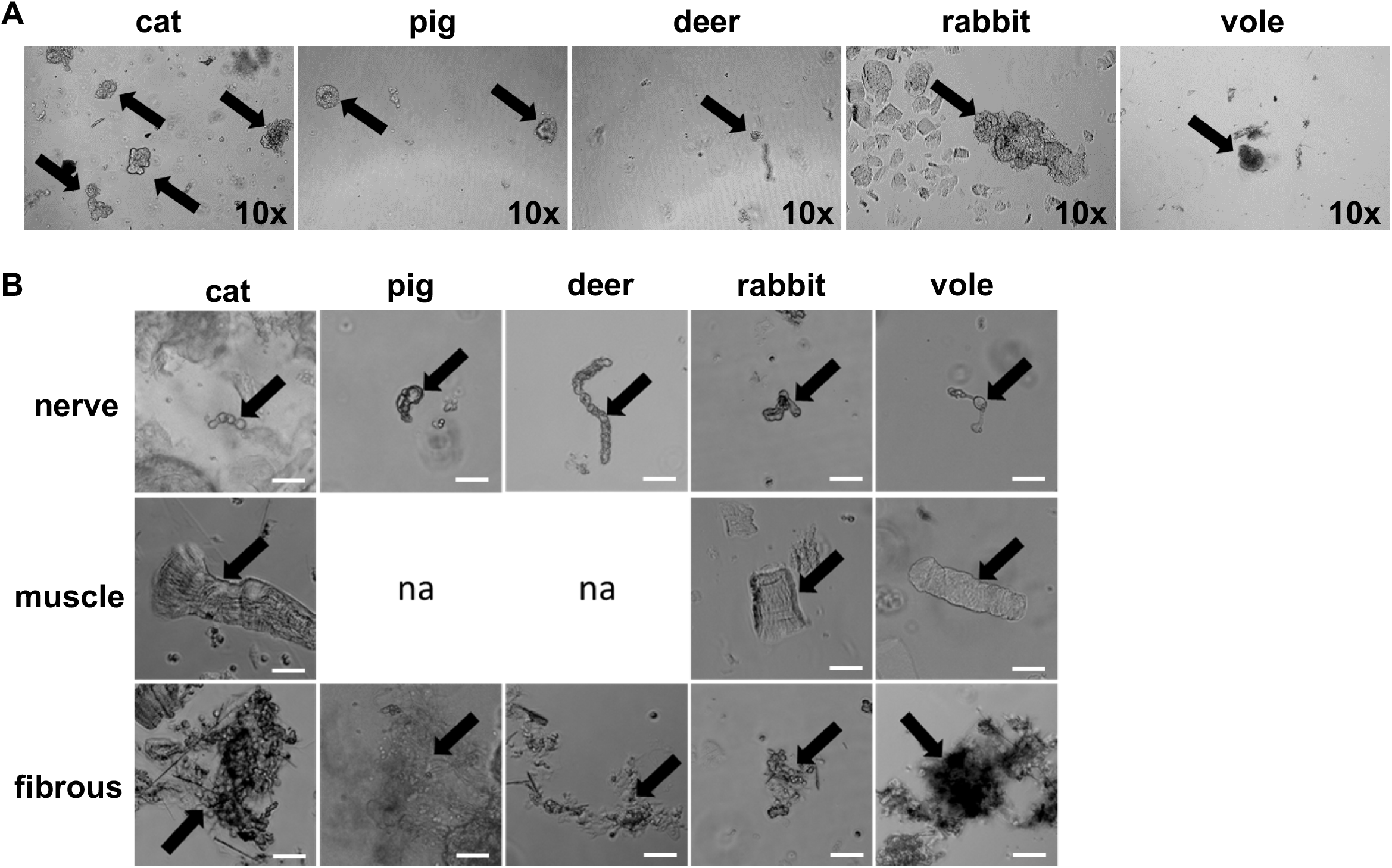
**(A)** Representative images (10X) of mammary tissue fragments (MTFs) from cat, pig, deer, rabbit, and vole, with fragments that will form bilayered organoids (black arrows). **(B)** Representative images of common contaminants seen from cat, pig, deer, rabbit, and vole MTF isolation including nerve tissue fragments (top row), muscle (middle row), and fibrous contamination (bottom row). Scale bars represent 50 μm.

**Fig. S6.**
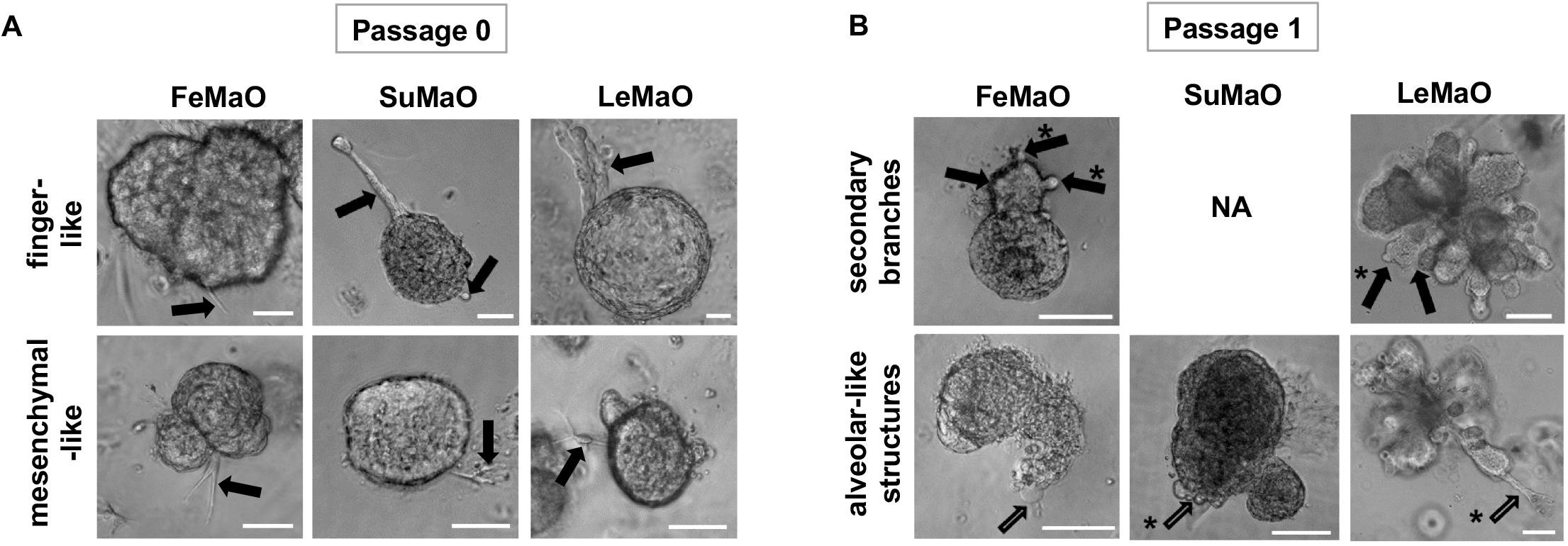
**(A)** Representative images of 3-day old FeMaOs, SuMaOs, and LeMaOs, at passage 0 with finger-like (top row) and mesenchymal-like (bottom row) protrusions, indicated by arrows. Scale bars represent 50 μm. (**B)** Representative images of 6-day old FeMaOs, SuMaOs, and LeMaOs, at passage 1. Primary buds and secondary branches are indicated by black arrows and black arrows with an asterisk, respectively (top row). Alveolar-like structures that are either bubble-like outgrowths or branch-like outgrowths are indicated by clear arrows and clear arrows with an asterisk, respectively. Scale bars represent 100 μm.

**Fig. S7.**
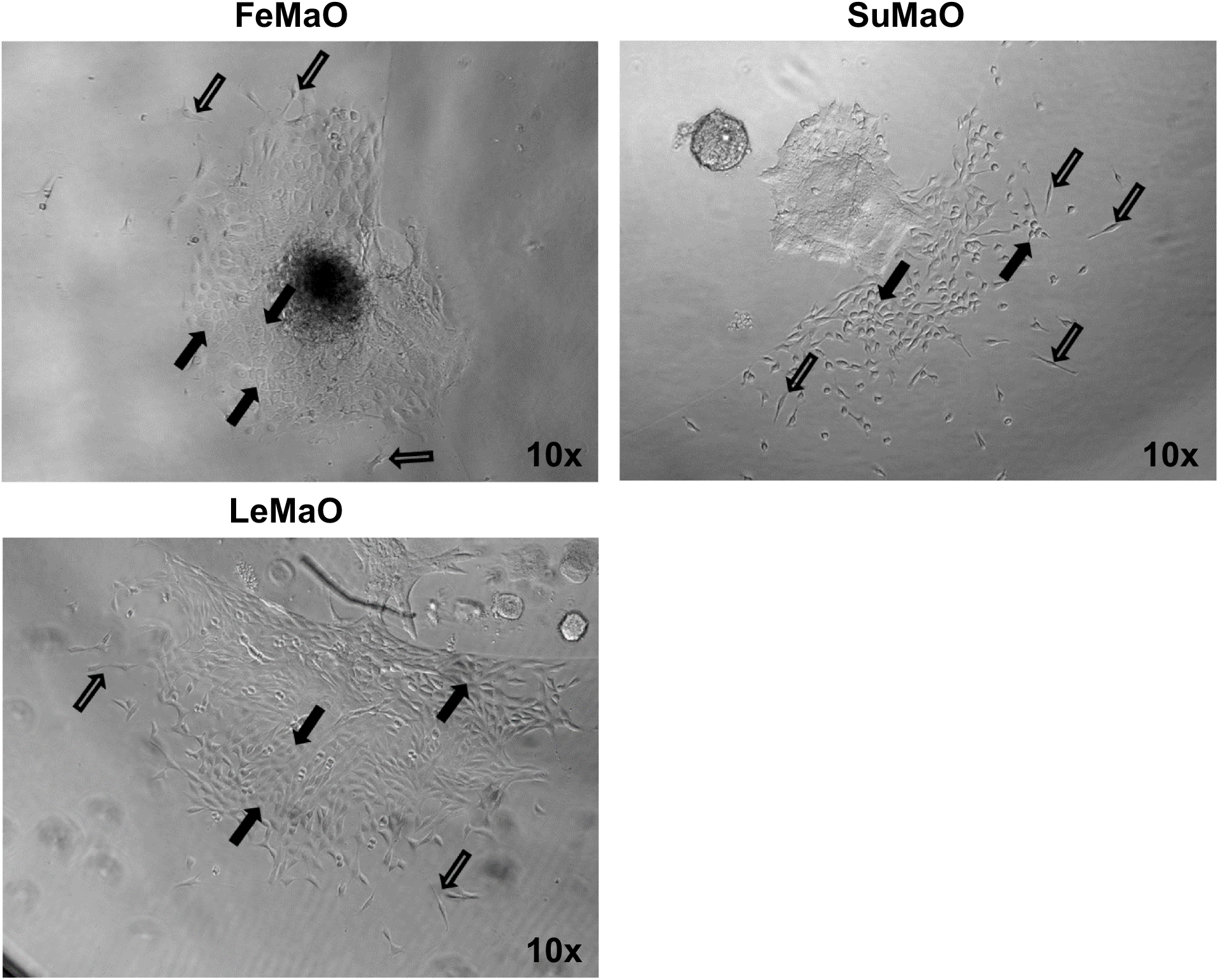
Representative image (10X) of 2D adherent cell cultures, after FeMaOs, SuMaOs, and LeMaOs, sank through the Matrigel to the bottom of the cell culture plates, containing cells with both mesenchymal-like morphology (clear arrows) and cobblestone morphology (black arrows).

